# Multicellular Model of Temozolomide Resistance in Glioblastoma Reveals Phenotypic Shifts in Drug Response and Migratory Potential

**DOI:** 10.1101/2025.07.08.663674

**Authors:** Victoria A. Kriuchkovskaia, Ela K. Eames, Sydney A. McKee, Paul J. Hergenrother, Rebecca B. Riggins, Brendan A.C. Harley

## Abstract

Glioblastoma (GBM) is the most common and aggressive primary malignant brain tumor in adults, with limited survival outcomes due to tumor recurrence, mainly driven by GBM cell invasion and therapy resistance. Although temozolomide (TMZ) remains the standard-of-care chemotherapeutic, its long-term efficacy is often compromised by rapid emergence of acquired resistance, largely mediated by the DNA repair enzyme, methylguanine methyltransferase (MGMT). To investigate the interplay between tumor heterogeneity, drug resistance, and the extracellular matrix (ECM) microenvironment, we adapted a 3D methacrylamide-functionalized gelatin (GelMA) hydrogel model to study the behavior of mixed populations of TMZ-sensitive and TMZ-resistant GBM cells. Using both single-cell distributions and multicellular spheroids, we report the impact of heterogeneous cell populations and TMZ dosing regimens, including physiological, supraphysiological, and metronomic TMZ schedules, on drug response and migration. We show that the combination therapy of TMZ with an MGMT inhibitor, lomeguatrib, can modulate TMZ resistance *in vitro*. This hydrogel model enables systematic investigation of GBM heterogeneity, “go-or-grow” phenotypic plasticity, and therapeutic resistance in an ECM-rich microenvironment, offering a valuable platform for future translational research.

## Introduction

Glioblastoma (GBM) is the most common and aggressive form of primary brain cancer in adults [1, 2]. Despite a multimodal treatment regimen that includes maximal safe surgical resection, radiation therapy (RT), and temozolomide (TMZ) chemotherapy, GBM patients show five-year survival rates of below 10% [3–5]. Even those who initially respond to TMZ often display rapid emergence of acquired resistance [6]. The complex tumor microenvironment (TME), a diverse cellular constituency that includes GBM stem cells, endothelial cells, astrocytes, microglia, and infiltrating macrophages as well as the extracellular matrix (ECM), can influence tumor progression. Beyond structural support, the ECM can modulate cancer cell growth, invasion, immune evasion, and therapeutic resistance, making it a potential driver of GBM progression [4, 7]. There is an opportunity to improve our collective understanding of acquired TMZ resistance in GBM via an approach that considers the contributions of the TME to GBM progression and therapeutic response in addition to conventional cancer-cell-intrinsicntinr sic drivers [8].

TMZ has remained a cornerstone of GBM therapy for over two decades, following the landmark clinical study by Stupp et al., which demonstrated that adding TMZ to RT significantly improved overall survival from 12.1 to 14.6 months, compared to RT alone [9, 10]. TMZ is an orally available, small-molecule chemotherapeutic that acts as an imidazotetrazine prodrug that undergoes intracellular hydrolysis to form the potent alkylating agent methyl diazonium [11, 12]. The cytotoxicity of TMZ is primarily due to DNA methylation with the O6-methylguanine (O6-MG) adduct the primary cause of the toxicity. In the cell, O6-MG mismatches with thymine, triggering the mismatch repair (MMR) system, where MMR incorrectly removes thymine from the undamaged strand, leading to further thymidine incorporation and ultimately apoptosis. However, O6-MG can also be repaired by the suicide DNA repair enzyme, methylguanine methyltransferase (MGMT), that is otherwise crucial for maintaining cellular genome stability [11, 14–16]. MGMT expression is regulated epigenetically, with methylation of its promoter leading to gene silencing. Hence, MGMT is a key GBM prognostic biomarker, and patients with silenced or low levels of MGMT generally show improved overall survival [13]. Unfortunately, TMZ treatment has been shown to induce MGMT expression in MGMT-low tumors via loss of methylation [14, 15], hence turning TMZ-sensitive tumors into ones that are TMZ resistant. There is an urgent need for new approaches to study and overcome MGMT-mediated TMZ resistance.

Tissue engineering platforms can be used to interrogate the role of the TME on GBM progression and drug efficacy [16–19]. While *in vivo* animal models remain the gold standard of pre-clinical research [17, 20], engineered models allow for systematic studies of cell migration, tumor growth, and metabolic activity [17, 21]. For instance, our lab has previously described a methacrylamide-functionalized gelatin (GelMA) hydrogel model of the GBM perivascular niche (PVN) to study the effect of angiocrine signals on GBM invasion, proliferation, and response to therapy, finding the PVN enhances GBM migration and reduces TMZ sensitivity [22–24]. More recently, we used hydrogel models to assess phenotypic shifts tied to TMZ resistance using isogenically-matched pairs of TMZ-sensitive and TMZ-resistant (via upregulated MGMT) GBM cell lines first generated by Tiek et al. [25] where long-term of the parental 42WT line to TMZ resulted in a TMZ-resistant line with increased MGMT expression and resistance to TMZ-induced apoptosis. Our prior work using these cells in 3D GelMA hydrogels established TMZ response curves to physiological, supraphysiological, and metronomic TMZ doses [26]. However, these and other studies have largely considered homogenous cell cultures (e.g., TMZ resistant cells vs. TMZ responsive cells only). As GBM tumors are not a cellular monolith but rather a complex and diverse multicellular entity, tissue engineering approaches provide the capacity to examine the behavior of heterogenous cohorts in a systematic fashion. Indeed, prior work in our laboratory using patient-derived GBM specimens showed stem-cell like subpopulations could induce a disparate response vs. the non-stem cell subpopulation in hydrogel models of the TME [27].

In this study, we describe the activity of mixed-cell cohorts in a GelMA-based 3D engineered model of the GBM TME (**Figure 1**). We report activity on mixtures of TMZ-sensitive and TMZ-resistant cell populations using both GBM cells distributed in GelMA hydrogels (to model low-density tumors) as well as spheroids embedded in GelMA hydrogels (to model high-density tumors). We report the role of the ratio of TMZ-sensitive vs. TMZ-resistant cell lines on TMZ response and cell motility across a wide range of TMZ doses, including physiologically relevant and metronomic dosing schedules. Further, we assess the response of multi-cellular cohorts to the combination of TMZ with an MGMT inhibitor, lomeguatrib (O6-BTG). This multi-cellular TMZ-responsive vs. TMZ-resistant model examines the emergence of “go-or-grow” phenotypic plasticity for GBM cohorts as well as its relation to acquired TMZ resistance.

**Figure 1.**
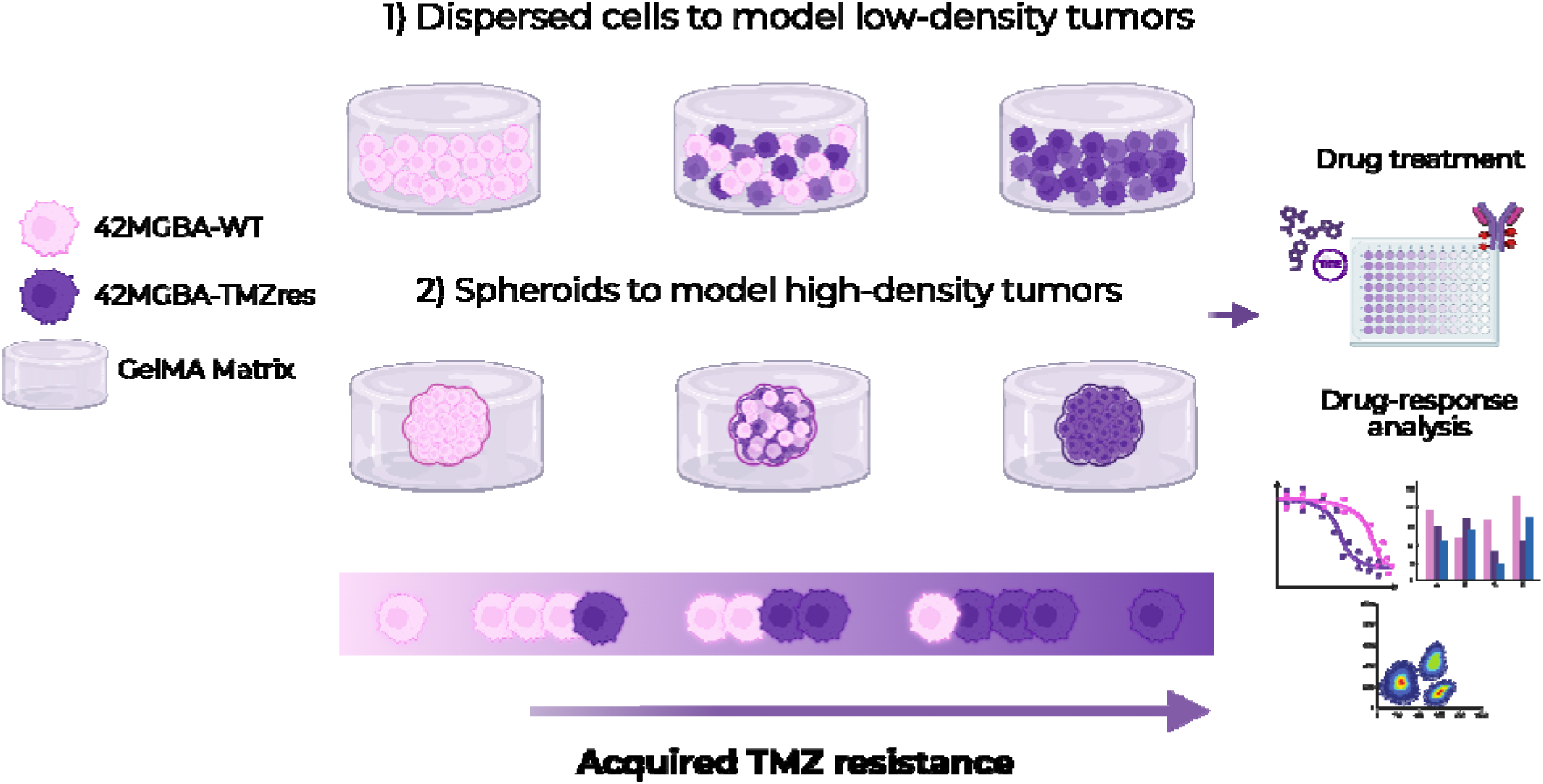
A multi-cellular model of glioblastoma (GBM) in a methacrylamide-functionalized gelatin (GelMA) hydrogel. Isogenically matched TMZ-sensitive and TMZ-resistant GBM cell lines (42MGBA-WT vs. 42MGBA-TMZres) are encapsulated as low-density tumor populations (distributed cells) or high-density tumor spheroids, to study cell motility and TMZ response. We report the effect of the ratio of sensitive to resistant cells on TMZ efficacy and migratory behavior via metabolic activity and spheroid-outgrowth assays.

## 2. Results

### 2.1. Co-Culturing TMZ-Sensitive and TMZ-Resistant Cell Lines in GelMA Hydrogels Creates a Continuum of Drug-Resistant Phenotypes

First, we generated a series of mixtures of temozolomide-sensitive (42WT) vs. TMZ-resistant (42TMZres) 42MGBA GBM cell lines distributed as individual cells within 20 μL GelMA hydrogels (10,000 total cells/sample) (**Figure 2A**): We studied responses for five different GBM mixtures: 100% 42WT + 0% 42TMZres (100WT); 75% 42WT + 25% 42TMZres (75WT); 50% 42WT + 50% 42TMZres (50WT); 25% 42WT + 75% 42TMZres (25WT); and 0% 42WT + 100% 42TMZres (0WT). We quantified drug response metrics **(Figure 2B and Figure S1)** based on the overall metabolic activity of each cell mixture in response to TMZ. Encapsulated GBM mixtures were treated with serial dilutions of TMZ, with the effect of single-dose TMZ treatments on cell viability assessed after 7 days of culture via alamarBlue HS™ assay. We generated growth-rate (GR) inhibition curves (**Figure 2B**) and the corresponding GR inhibition metric (**Figure S1**) for TMZ using methods described by Hafner et al. [28] and our own prior work [29]. The GR50 values varied from 6.6 μM (100WT) to >1300 μM for (0WT), with GR50 values for cell mixtures staying within the bounds established by the homogeneous 42WT and 42TMZres only controls: 459 μM (75WT); 881 μM (50WT); 1094 μM (25WT).

**Figure 2.**
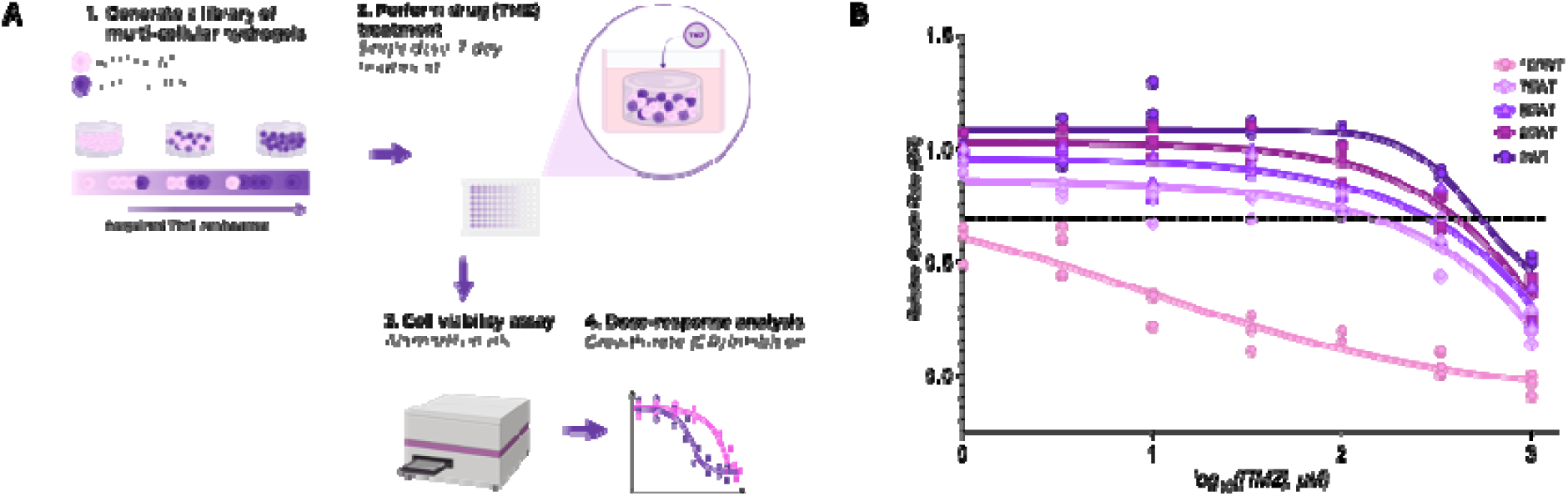
Co-culturing TMZ-sensitive and TMZ-resistant cell lines in GelMA hydrogels creates a library of multicellular hydrogels with distinct drug-resistant niches. **A**. Schematic illustrating the generation of a library of multicellular GelMA hydrogel models to investigate TMZ response of mixtures of TMZ-resistant vs. TMZ-responsive GBM cell lines. **B**. Growth-rate (GR) inhibition curves for each cellular blend based on the metabolic activity 7 days post treatment in response to a single-dose TMZ administration. Each GR value is shown (n = 3). GR curves are shown as fitted model described in Hafner et al.[28] using the Online GR Calculator (www.grcalculator.org/grcalculator) from n = 3 replicates per TMZ concentration. Figures were created in BioRender.com and GraphPad Prism.

### 2.2. The Migratory Capacity of Drug-Resistant GBM Spheroids are Sensitive to Single-Dose TMZ Treatment

We subsequently evaluated the migratory capacity of multi-cell cohorts after single-dose TMZ treatments. We embedded GBM spheroids (5,000 total cells/spheroid) in 20 μL GelMA hydrogels; **Figure 3A**) composed of the same ratios (by cell number) of 42WT and 42TMZres GBM cell lines (100WT; 75WT; 50WT; 25WT; 0WT). After 24 hours of culture, spheroids were treated with single TMZ doses of 0 μM (DMSO control), 20 μM (physiologically relevant), 100 μM (high), or 1000 μM (supra-physiological). Cell viability was quantified after 7 days via an alamarBlue HS™ assay (**Figure 3B**). Unlike dispersed cultures, a single-dose TMZ treatment did not induce significant changes in metabolic activity of multicellular spheroids (p = 0.9501); the interaction effect between TMZ dose and cell mixtures was also not significant (p = 0.8928). However, the radial migration of cells from GBM spheroids 7 days after TMZ treatment (**Figure 3C and Figure S1**) was significantly reduced in response to single-dose TMZ treatment (**Figure 3C**). Single-dose TMZ treatment (p < 0.0001) and the cell-mix composition (p < 0.0001) both significantly influenced cell migration (though the interaction between these two factors was not significant p = 0.0798, at the risk of a Type I error of 0.05).

**Figure 3.**
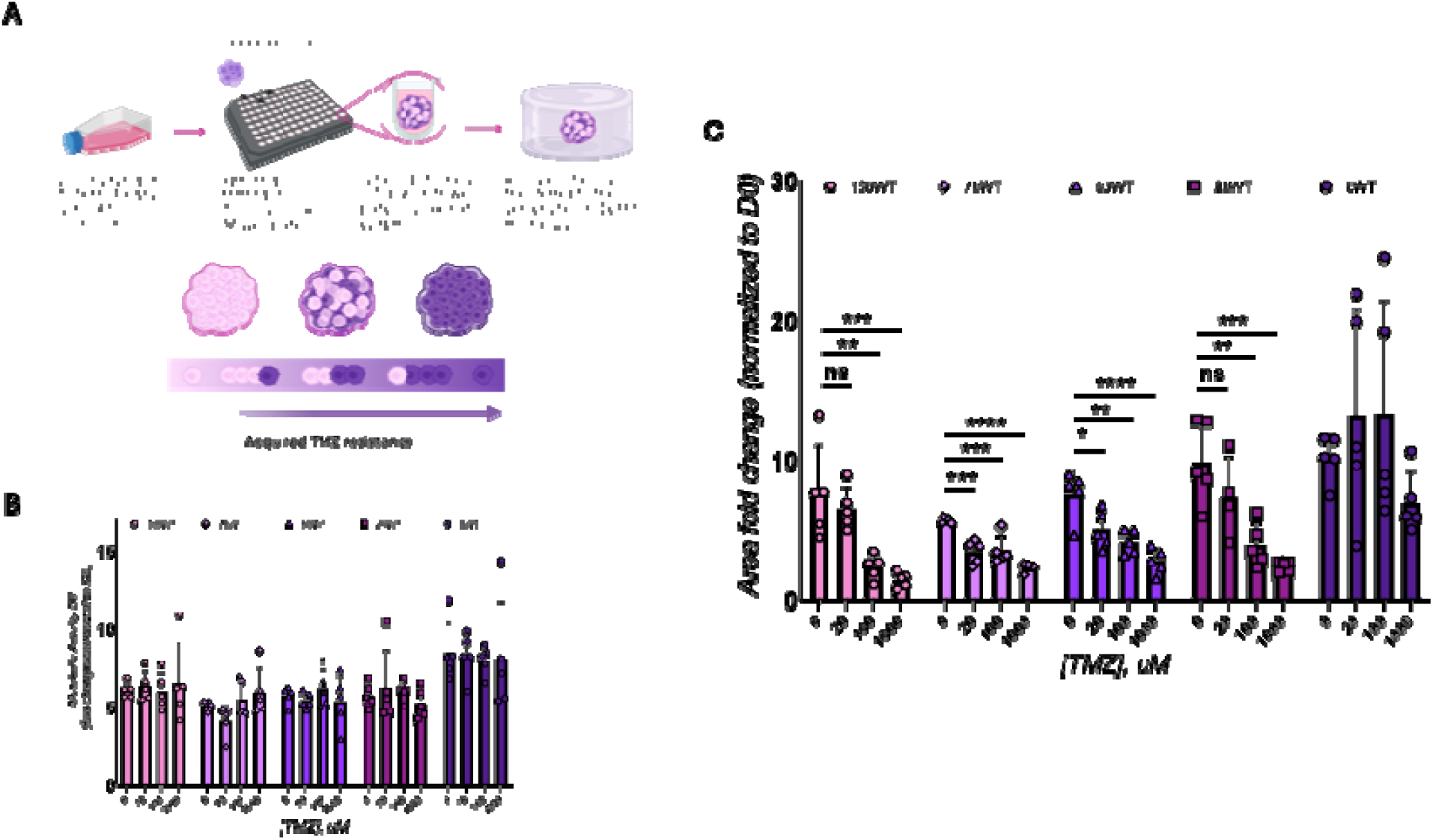
The migratory and proliferative behaviors of drug-resistant GBM cell niches after single-dose TMZ treatment. **A**. Schematic illustrating analysis of 5,000 cell spheroids composed of mixtures of TMZ-sensitive (42WT) and TMZ-resistant (42TMZres) GBM cells. **B**. The metabolic activity after single-dose TMZ treatment at concentrations of 0 µM (DMSO control), 20 µM (physiologically relevant), 100 µM (high), and 1000 µM (supra-physiological) measured 7 days post-treatment. **C**. GBM migratory capacity as measured as fold change of the total area of spheroid outgrowth vs. day 0 spheroid area revealed significant effects of single-dose TMZ treatment. *: p < 0.05. **: p < 0.01. ***: p < 0.001. ****: p < 0.0001. Figures were created in BioRender.com and GraphPad Prism.

### 2.3 Metronomic TMZ Dosing of GBM Spheroids Exhibit “Go-or-Grow” Phenotype

We then investigated the migratory and proliferative capacity of GBM spheroids (5,000 total cells/spheroids embedded in 20 μL GelMA hydrogels) containing mixtures of 42WT and 42TMZres cells treated with 5 daily TMZ doses of either 0 μM TMZ (DMSO control) or 20 μM TMZ. The metabolic activity (alamarBlue HS™ assay 7 days after the final treatment) of all groups containing TMZ response cells (100WT, 75WT, 50WT, 25WT) was significantly reduced with metronomic dosing (**Figure 4A**). Metronomic dosing (p < 0.0001) and starting cell-mix composition (p < 0.0001) both influenced metabolic activity; there was also a significant interaction between these factors (p = 0.0023). Radial patterns of cell migration (**Figure 4B and Figure S2**) into the GelMA hydrogel were also evaluated 7 days after the final TMZ treatment (**Figure 4B**). Interestingly, spheroids containing high fractions of TMZ-responsive cells showed reduced radial motility compared to spheroids comprise of high fractions of TMZ resistant cells. While statistical analysis revealed that metronomic TMZ treatment did not significantly affect overall migration (p = 0.5263), the cell-mix composition (p < 0.0001) as well as the interaction between these factors (p < 0.0001) both significantly affected invasion. Broadly, metronomic TMZ treatment led to a reduction in in metabolic activity and migratory capacity for most groups. However, the 50WT group displayed a different phenotype, with reduced metabolic activity but significantly enhanced migratory phenotype after exposure to metronomic TMZ treatment at a physiological dosing regimen (**Figure 4A and Figure 4B**; 5 x 20 μM TMZ).

**Figure 4.**
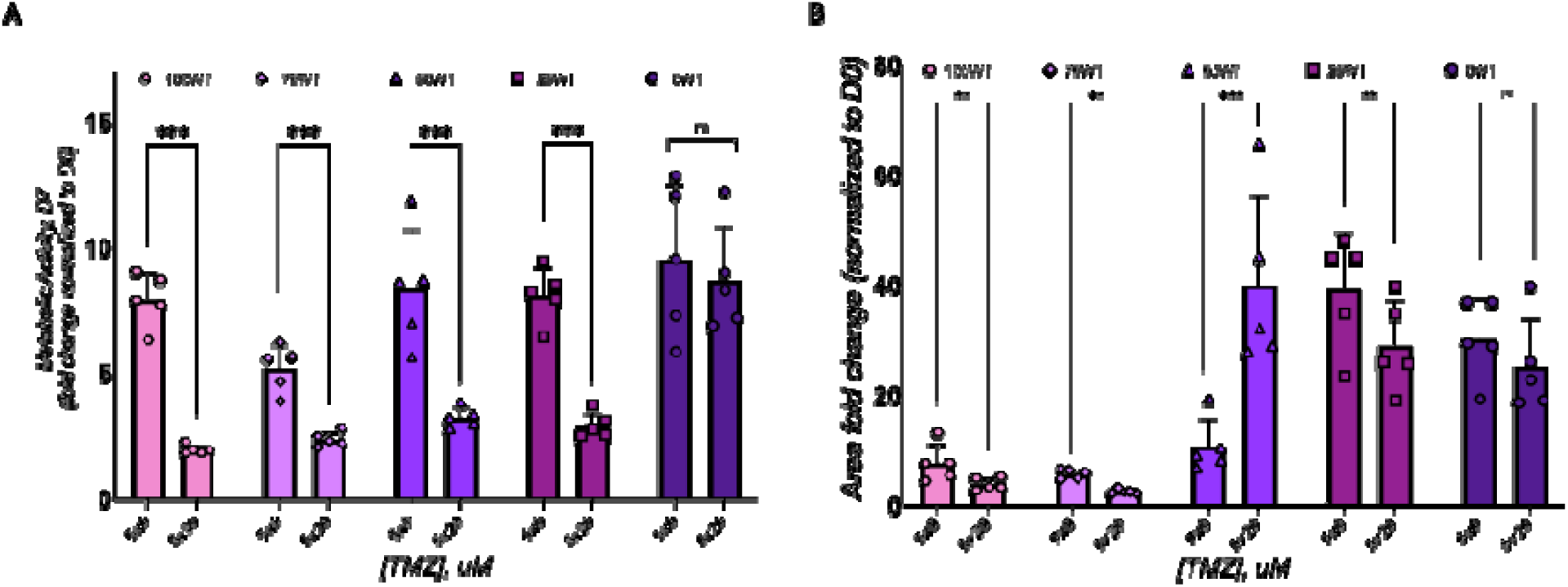
Metronomic TMZ schedule administered to GBM spheroids in GelMA hydrogels. Spheroids composed of various ratios of TMZ-sensitive (42WT) and TMZ-resistant (42TMZres) GBM cell lines were encapsulated in GelMA hydrogels (5,000 total cells per spheroid in 20 µL hydrogels). After 1 day of culture, spheroids were treated with five daily doses of TMZ (20 µM) or DMSO control (0 µM). **A**. The metabolic activity measured using the alamarBlue HS™ assay 7 days after the final TMZ treatment. **B**. The migratory capacity of spheroids measured using a spheroid-based invasion assay 7 days after the final TMZ treatment. While most groups showed reduced metabolic activity and migratory capacity in response to metronomic TMZ, the 50% 42WT + 50% 42TMZres group exhibited significantly increased migratory behavior compared to DMSO control, suggesting a unique response to metronomic TMZ treatment. Data are shown as individual data points, mean, and SD. *: p < 0.05. **: p < 0.01. ***: p < 0.001. Figures were created in GraphPad Prism.

### 2.4. The Combination of TMZ with an MGMT Inhibitor, Lomeguatrib, Enhances the Effect of Single-Dose TMZ Treatment on GBM Metabolic Activity

Finally, we examined the efficacy of the combination of a single-dose TMZ treatment with 4-hour pre-treatment with an MGMT inhibitor, lomeguatrib, on the metabolic activity of dispersed populations of TMZ resistant and WT GBM cells. Here, we focused on a subset of mixtures (0WT; 50WT; 100WT) and treated each with the GR50 dose calculated for each GBM cell mixture (**Figure 2**); this allowed us to effectively compare the combined efficacy of TMZ and the MGMT inhibitor across groups but required us to only use the distributed cells cultures that were used to calculate GR50 dosages. Dispersed GBM cohorts were encapsulated into GelMA hydrogels for 24 hours then treated with lomeguatrib (50 µM) or a DMSO control for 4 hours. Each group was subsequently treated with a single dose of TMZ at their unique GR50 dose (100WT: 6.6 µM; 50WT: 881 µM; 0WT: 1094 µM) or with a DMSO control as a way to compare relative efficacy of MGMTi+TMZ treatments. MGMTi exposure alone did not influence the metabolic activity of TMZ-sensitive (100WT) cells (**Figure 5A**) or the 50WT (50WT/50TMZres) mixed groups (**Figure 5B**). However, MGMTi alone significantly reduced the metabolic activity of the fully TMZ-resistant (0WT) group (**Figure 5C**). Moreover, the combination of TMZ (at the calculated GR50 for each group) and MGMTi significantly reduced the metabolic activity of experimental groups containing TMZ-resistant cell populations for all cohorts that contained TMZ-resistant cells (**Figure 5B-C**), but not for the cohort containing only WT cells (**Figure 5A**; 100WT). Consistent with this, median-effect analysis shows that the MGMTi+TMZ combination is synergistic in both the 50WT (relative index, RI = 2.4) and 0WT groups (RI = 1.4).

**Figure 5.**
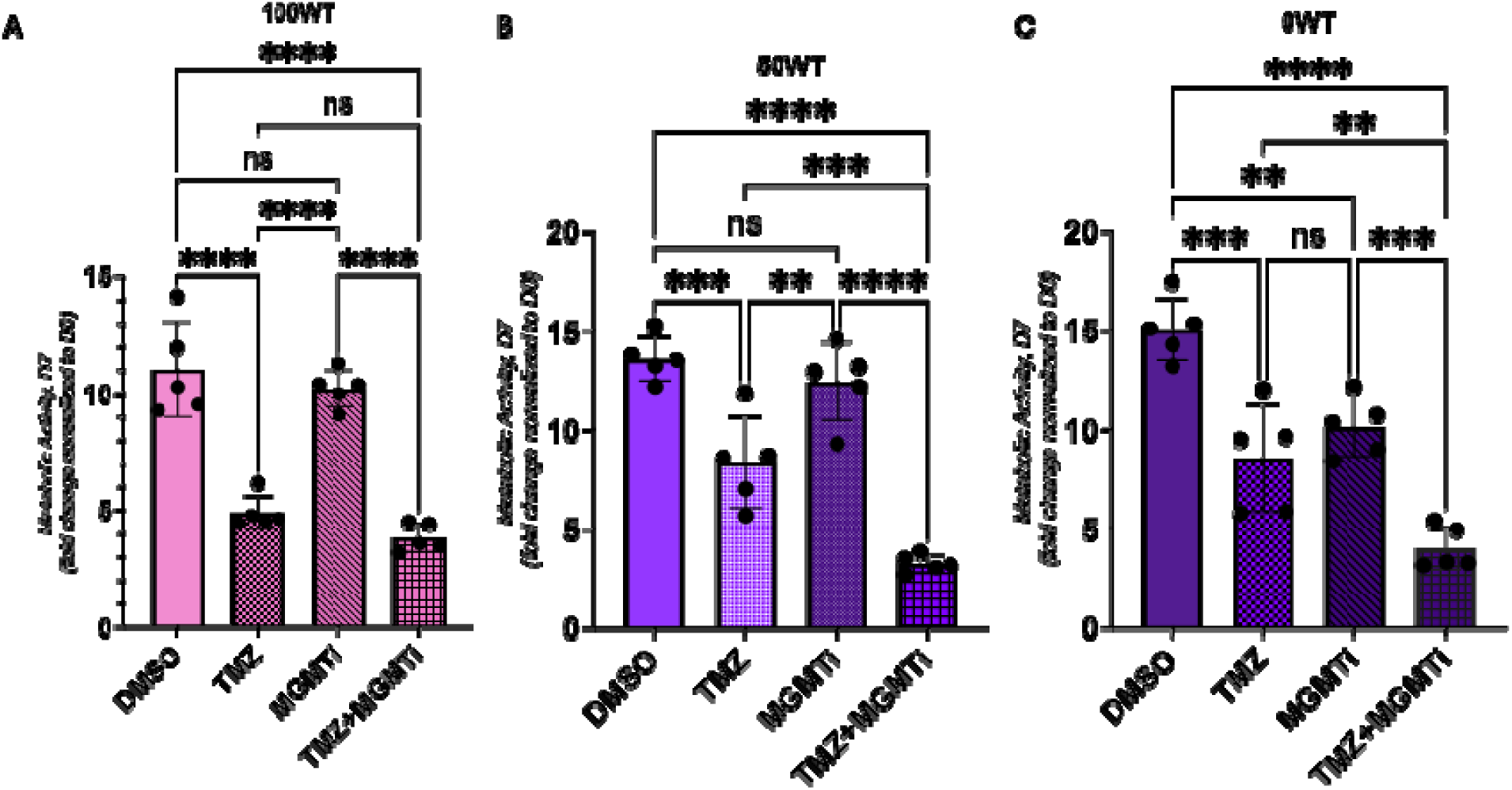
Combination treatment of TMZ with an MGMT inhibitor, lomeguatrib, enhances the effect of a single-dose TMZ treatment on multicellular GBM cultures. **A**. Effect of 4-hour pre-treatment with lomeguatrib (50 µM) on the metabolic activity of the control group containing only TMZ-sensitive cells (100WT) following single-dose TMZ treatment (6.6 µM). No significant effect of MGMTi pre-treatment was observed on the metabolic activity in this group . **B**. Impact of lomeguatrib on the 50WT group after single-dose TMZ treatment (881 µM). No statistically significant effect was observed with MGMTi pre-treatment alone, while combination treatment was more effective compared to TMZ alone. **C**. Metabolic activity of the fully TMZ-resistant group (0WT) following MGMTi pre-treatment and single-dose TMZ treatment (1094 µM). MGMTi alone significantly reduced the metabolic activity in the 0WT group. Combination treatment was more effective than TMZ alone. Data are shown as individual data points, mean, and SD. *: p < 0.05. **: p < 0.01. ***: p < 0.001 (one-way ANOVA with Tukey’s). Figures were created in GraphPad Prism.

## 3. Discussion

In this project, we describe a hydrogel system to evaluate the phenotype of heterogeneous mixtures of TMZ resistant and TMZ responsive cells in response to TMZ therapy. The GBM TME is complex and requires tools to study heterogeneous cell-matrix and cell-cell interactions that vary spatially and with time [4, 7]. TMZ may significantly affect both cell proliferation/viability as well as their invasive capacity [8, 30]. The “go-or-grow” hypothesis suggests GBM can reversibly switch between migratory and proliferative phenotypic states [31, 32]. Originally proposed to explain the mutual exclusivity of migration and proliferation in astrocytoma cells, *in vivo* evidence has suggested proliferation may be lower at the tumor perimeter and higher in the core (migratory cells may be less proliferative) [33]. More recently, the concept of go-or-grow plasticity suggests single GBM cells may reversibly switch between the proliferative and invasive phenotypes to avoid therapies targeting a single cellular phenotypic state [31, 34–36]. This switching is likely complicated by the cellular heterogeneity of the GBM tumor, which may include mixtures of TMZ-resistant and responsive cells, whose relative population distribution is likely to evolve in time in response to TMZ treatment. While challenging to study *in vivo*, engineered models of the TME provide a unique opportunity to evaluate the behavior of populations of GBM cells that exhibit disparate levels of TMZ response.

The 42WT line and its isogenically-matched 42TMZres line offer an ideal cellular system to study “go-or-grow” phenotypes. Originally described by Tiek et al. [25], long-term exposure of the parental 42WT line to TMZ resulted in a TMZ-resistant line with increased MGMT expression and resistance to TMZ-induced apoptosis. More recently we observed changes in TMZ sensitivity, cell invasion, and matrix-remodeling cytokine production for homogenous populations of TMZ-resistant vs. TMZ responsive (wild type) cells. We investigated the behavior of TMZ-sensitive and TMZ-resistant GBM cell dispersed in the GelMA hydrogel to model low-density tumors with actively dividing cells (thus modeling the “grow” sub-state). We then embedded GBM spheroids (5,000 cells/spheroid) into the GBM hydrogel to mimic high-density tumors more likely to exhibit substantial diffusive infiltration (the “go” sub-state). Interestingly, while spheroid models offer a model to study the migratory GBM phenotype, cells exhibit reduced proliferation which led to reduced efficacy of TMZ on the GBM spheroids vs. distributed GBM cells, especially at physiologically relevant doses. We subsequently quantified migratory and proliferative capacity of GBM spheroids in response to metronomic TMZ dosing (repeated 20 µM dose). Metronomic TMZ dosing reduced the metabolic activity of the spheroids containing drug-sensitive (WT) subpopulations and reduced the invasive capacity of GBM spheroids containing primarily TMZ-sensitive (100TW, 75WT) GBM cells. Interestingly, the 50WT/50TMZres (50WT) cohort displayed a significant increase in migratory capacity but significant decrease in metabolic activity in response to metronomic 20 µM TMZ doses. These findings suggest future use of GBM spheroid models comprised of an initially equal mix of WT and TMZres cells may be particularly valuable to study the de-coupled effect of TMZ treatment on the “go-or-grow’ phenomena. Such a model would be ideally suited for the use of selective labeling tools to better assess the size and influence of TMZ resistant versus wild type subpopulation and would be valuable precursors to future use of patient-derived specimens that contain heterogenous mixtures of stem-like cell populations previously shown to influence cell invasion [27].

Recently, there has been an increased interest in studying the combination of MGMT inhibitors and O6-alkylating chemotherapeutics to increase the cytotoxicity of alkylating agents against tumor cells [37–40]. The 42TMZres cell line used in our model has previously been shown to exhibit robust MGMT expression (vs. none in the 42WT) [41], motivating analysis of the combined efficacy of a single-dose TMZ treatment after pre-treatment with lomeguatrib. Excitingly, groups containing TMZ-resistant cells (50WT; 0WT) showed increased efficacy of TMZ treatment with lomeguatrib. Although lomeguatrib alone did not reduce the metabolic activity of the 100WT or 50WT groups, it significantly reduced the efficacy of the homogenous TMZ-resistant (0WT) group. While lomeguatrib was administered at a concentration reported to be non-toxic to GBM cells in vitro (50 µM) [37, 40], the significant inhibitory effect of lomeguatrib alone after 4-hours exposure on 0WT GBM group may due to the degree of MGMT overexpression in the 42TMZres cells [41]. Indeed, lomeguatrib is a highly specific and potent MGMT inhibitor previously shown to significantly reduce MGMT protein levels in vitro at even low doses (0.01 µM) in human GBM cell lines with unmethylated MGMT promoter [40]. Taspinar et al. [42] and Ugur et al. [43] showed that combination of TMZ and lomeguatrib (at 50 μM) administered to GBM and anaplastic astrocytoma cell lines reduced TMZ IC50, decreased

MGMT expression, and increased p53 expression without observable changes in cell cycle, suggesting apoptosis caused by increased p53 expression and decreased MGMT expression. Expanded consideration of the pharmacokinetics of MGMT inhibitors within the tumor microenvironment would significantly improve future studies modeling the combination of TMZ and MGMT inhibitors *in vitro*. Already, ongoing work is determining changes in GBM invasion in response to combinations to metronomic TMZ and limiting dilutions of lomeguatrib to establish more complex shifts in GBM phenotype with combination therapy.

## 4. Conclusions

This project describes the use of a GelMA hydrogel model containing distributed GBM cells or dense GBM spheroids to investigate aspects of “go-or-grow” plasticity in response to acquired MGMT-mediated TMZ resistance of GBM. We identify the limited efficacy of single-dose TMZ treatments on metabolic activity of TMZ-resistant dense tumor models, show single low-dose TMZ treatments can impact the migratory capacity of TMZ-sensitive and TMZ-resistant GBM populations, and demonstrate a 50:50 blend of TMZ responsive and resistant cells may have particular value as an *in vitro* model of the go-or-grow plasticity of GBM cells. Here, heterogenous mixtures of TMZ-responsive and TMZ-resistant GBM cells to metronomic TMZ doses reveals a migration-proliferation dichotomy. We also show that heterogenous mixtures of TMZ-responsive and TMZ-resistant GBM cells can be used to assess the efficacy of combination treatment with TMZ and a common MGMT inhibitor, lomeguatrib. Together, these findings establish the foundation for future efforts to exploring the mechanisms underlying the “go-or-grow” plasticity using well-characterized cell line or patient-derived cell specimens. Such studies are important steps towards identifying new treatment regimens to overcoming the challenges posed by MGMT-mediated acquired TMZ resistance in GBM.

## 5. Experimental Section

### 5.1. 2D Cell Culture

The isogenically-matched 42MGBA-WT (42WT) and 42MGBA-TMZres (42TMZres) cells were provided by Dr. Rebecca B. Riggins (Lombardi Comprehensive Cancer Center (LCCC), Georgetown University, Washington, D.C.). All cells tested negative for Mycoplasma contamination using MycoStrip™ - Mycoplasma Detection Kit (InvivoGen, San Diego, CA) and by PCR test done at The Tumor Engineering and Phenotyping (TEP) Shared Resource, University of Illinois Urbana-Champaign, Urbana, IL. Cells were grown in adherent culture flasks in DMEM (Cat. #11965, Thermo Fisher Scientific, Waltham, MA) supplemented with 10% FBS (R&D Systems, Minneapolis, MN) and 0.2% v/v plasmocin (InvivoGen, San Diego, CA). All cells were passaged fewer than 10 times and maintained in a humidified incubator with 95% air and 5% CO2 at 37 °C.

### 5.2. Compound Preparation

Temozolomide (TMZ; Cat. #S1237, Selleckchem, Houston, TX) was dissolved in DMSO (Thermo Fisher Scientific, Waltham, MA) to a stock concentration of 130_mM. For all experiments, the dissolved TMZ stock was subsequently added to growth media to achieve the desired TMZ concentration immediately before supplementing TMZ-containing media to cells. Lomeguatrib (Cat. #11732, Cayman Chemical, Ann Arbor, MI) was dissolved in DMSO (Thermo Fisher Scientific, Waltham, MA) to a stock concentration of 15_mM. For the experiments, lomeguatrib stock solution was subsequently added to growth media at the desired concentration (50 μM) immediately before adding lomeguatrib-containing media to cells.

### 5.3. Methacrylamide-Functionalized Gelatin (GelMA) Synthesis and Characterization

GelMA macromer (hydrogel precursor) was synthesized as previously described. Briefly, 1 g of porcine gelatin type A, 300 bloom (Sigma Aldrich, St. Louis, MO) was dissolved in 10 mL of carbonate-bicarbonate (CB) buffer (pH 9.4) at 50 °C. Subsequently, 40 µL of methacrylic anhydride (Sigma Aldrich, St. Louis, MO) was added dropwise, and the reaction proceeded for 1 hour with vigorous stirring (400 RPM). The reaction was quenched with 40 mL of warm deionized water and dialyzed in 12-14 kDa dialysis membranes for 7 days against deionized water with daily water exchange. The product was then frozen and lyophilized. 1HNMR was used to determine the degree of functionalization (DOF). GelMA with DOF of ∼55% was used in this study. Compressive moduli of GelMA hydrogels were measured using an Instron 5943 mechanical tester (Norwood, MA). Hydrogels were tested under unconfined compression at the rate of 0.1 mm/min, with the Young’s modulus obtained from the linear region of the stress-strain curve (2.5–17.5% strain) using a custom MATLAB (MathWorks, Natick, MA) code.

### 5.4. Fabrication of Low-Density Dispersed Cell-Laden GelMA Hydrogels

Glioblastoma cells of interest (42WT or 42TMZres) were homogeneously resuspended in the GelMA precursor solution at the concentration of 5 x 10^5^ cells/mL, and the resulting cell suspension was pipetted into custom Teflon molds (5 mm diameter, 1 mm thick). Hydrogels were formed after photopolymerization for 45 seconds using a UV lamp (λ = 365 nm, 5.69 mW/cm2). Hydrogels were deposited into 48-well plates with each well containing 500 µL of growth media. Hydrogels were further cultured in a humidified incubator with 95% air and 5% CO_2_ at 37 °C for subsequent experiments.

### 5.5. Metabolic Activity and Viability Assay

Cell viability and metabolic activity were measured with alamarBlue™ HS Cell Viability Reagent (Thermo Fisher Scientific, Waltham, MA) following the manufacturer’s protocol. Briefly, cells were encapsulated into GelMA hydrogels as described above. At the time point of interest, growth medium was aspirated from the wells and new growth medium (450 µL) was added to the hydrogels, and the alamarBlue™ solution (10% final volume/50 µL) was added to each well. After 2-hour incubation on a shaker (60 rpm) in a humidified incubator with 95% air and 5% CO_2_ at 37 °C, the alamarBlue™ solution was measured for the fluorescence of resorufin (540 (52)-nm excitation, 580 (20-nm emission) using a F200 spectrophotometer (Tecan, Switzerland). Negative control (background) was subtracted from each measurement.

### 5.6. Drug Response: Growth-Rate (GR) Inhibition Assay

Drug response metrics were based on cell viability/metabolic activity assay and measured with alamarBlue™ as described above. To generate drug response curves, each of the cell lines (42WT or 42TMZres) were encapsulated into GelMA hydrogels at 5 x 10^5^ cells/mL as described above. After 24 hours of culture, initial viability (D0) was determined with alamarBlue assay as described above. After D0 measurement were collected, hydrogels were rinsed with PBS, and growth medium was supplemented with the drug of interest (e.g., TMZ) and added to the hydrogel culture as single-dose treatments ranging from 0.03 µM to 1000_µM (serial dilutions) or a DMSO vehicle control (0_μM). Without replenishing the drug, drug response was measured 7 days after drug treatment using alamarBlue™ assay. Growth rate (GR) inhibition and related drug response metrics (GR50) were determined, as described in Hafner et al. First, relative cell count was determined based on the metabolic activity measured with the alamarBlue™ assay. Then, GR metrics were calculated as described by Hafner et al., and the drug response metrics were calculated using the Online GR Calculator (www.grcalculator.org/grcalculator).

### 5.7. Metronomic Drug Schedule Assay

To evaluate the effect of metronomic (low-dose, continuous) TMZ exposure, cell-laden GelMA hydrogels were treated with a single dose of 20 µM, or 5 repeated doses of 20 µM, or a cumulative single dose of 100 µM. The repeated doses were administered in daily intervals (24 hours between each dose). Cell viability assay (as described above) was done at days 0, 3, 5, and 7 after either the single TMZ treatment for single dose exposures or after the final TMZ treatment for continual, metronomic dose exposure. Day 0 was set to be as the day of the final TMZ treatment for each experiment. Data was normalized to day 0 measurements prior to the analysis.

### 5.8. MGMT Inhibitor Combination Treatment Assay

GBM cells were first incapsulated into GelMA-based hydrogels using the dispersed cell model described above. After 24 hours in the hydrogel culture, cells were pretreated with lomeguatrib (Cat. #11732, Cayman Chemical, Ann Arbor, MI) for 4 hours. Following the lomeguatrib treatment, cells were washed with phosphate-buffered saline (PBS) to remove any residual compound. The cells were then treated with the selected compound for an additional 7 days.

### 5.9. Spheroid Migration Assay

Spheroids were constructed using the previously reported method [26]. Briefly, GBM cells (42WT/42TMZres) were counted and resuspended into 5000 cells/200 μL media per well and added into 96-well spheroid ultra-low attachment (ULA) microplates (Corning, Corning, NY). Plates were centrifuged at 100×g for 1 min to assist spheroid formation then placed into a humidified incubator (37°C, 5% CO2) for 24 h. Plates were then incubated for additional 24 h with gentle shaking at 60 rpm. Subsequently, formed spheroids were transferred and mixed with GelMA precursor solution and photopolymerized into hydrogels as described above. Spheroid images were acquired using a Leica DMI8 bright-field microscope (Leica, Germany) at days 0 (immediately upon seeding), 1, 2 and 3. Invasion was then quantified via ImageJ, and the invasion distance was reported as fold change of the spheroid outgrowth area compared to day 0 as described previously.

### 5.10. Statistical Analysis

All statistical analyses were performed using GraphPad Prism (version 10.4.1). Additionally, all graphs were generated in GraphPad Prism. Data reported in this manuscript were plotted as the mean ± standard deviation (SD). Each experimental condition was performed in 3-5 replicates to conduct statistical analysis. Normality was determined using the Shapiro–Wilk test and homoscedasticity was determined *via* Bartlett’s test for normal data or Levene’s test for non-normal data. Comparisons between two unpaired groups were performed using a Mann-Whitney U-test, while comparisons between multiple groups were performed using a two-way ANOVA when assumptions were met with cellular-mix composition and TMZ dosing being used as the two factors. Tukey’s post-hoc test was used to compare significance between these groups. The level of significance, α, representing the chance of making a Type I Error, was set to 0.05 for all analyses. For synergy analysis, median-effect analysis was used to calculate the Relative Index (RI) as in [44, 45]. RI values were obtained by calculating the expected cell metabolic activity (A_exp_; the product of activity obtained with drug A alone and the activity obtained with drug B alone) and dividing A_exp_ by the observed cell metabolic activity in the presence of both drugs (A_obs_). A_exp_/A_obs_ > 1.0 indicates a synergistic interaction.

## Acknowledgements

The content herein is solely the responsibility of the authors and does not necessarily represent the official views of the National Institutes of Health. The authors also gratefully acknowledge additional funding provided by the Department of Chemical & Biomolecular Engineering, The Department of Chemistry, and Cancer Center at Illinois at the University of Illinois at Urbana-Champaign.

## Author contributions

We describe contributions to the manuscript using the Contributor Roles Taxonomy (CRediT) [46, 47]:

Conceptualization: VK, SAM, PJH, RBR, BACH Methodology: VK

Investigation: VK, EE Visualization: VK

Formal Analysis: VK, RBR Data Curation: VK

Project Administration: BACH Supervision: BACH, RBR Writing—original draft: VK

Writing—review & editing: VK, PJH, RBR, BACH Resources: BACH

Funding Acquisition: RBR, PJH, BACH

