## Supplemental Information for "Multicellular Model of Temozolomide Resistance in Glioblastoma Reveals Phenotypic Shifts in Drug Response and Migratory Potential"

^1^ Dept. Chemical and Biomolecular Engineering

^2^ Carl R. Woese Institute for Genomic Biology

^3^ Cancer Center at Illinois

^4^ Dept. of Chemistry

University of Illinois Urbana-Champaign

Urbana, IL 61801, USA.

^5^ Dept. of Oncology, Lombardi Comprehensive Cancer Center,

Georgetown University Medical Center

Washington, DC 20057, USA.

**Corresponding Author:**

B.A.C. Harley

Dept. of Chemical and Biomolecular Engineering

Cancer Center at Illinois

Carl R. Woese Institute for Genomic Biology

University of Illinois at Urbana-Champaign

110 Roger Adams Laboratory

600 S. Mathews Ave.

Urbana, IL 61801

**Supplemental Figures**

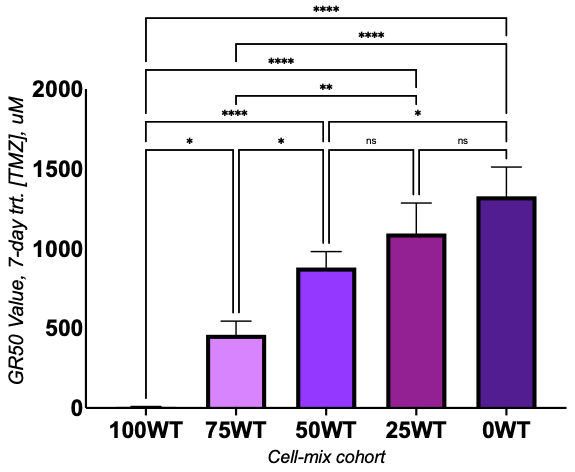

**Figure S1. Calculated TMZ GR50 values for each group 7-days post single-dose TMZ treatment.** GR values showed a progressive increase in resistance from the 42WT-only group (6.6 µM) to the 42TMZres-only group (1327 µM). GR50 values for mixed cell populations were intermediate, with the 75WT group at 450 µM, 50WT at 881 µM, and 25WT at 1094 µM. *: p < 0.05. **: p < 0.01. ***: p < 0.001. Figure was created in GraphPad Prism.

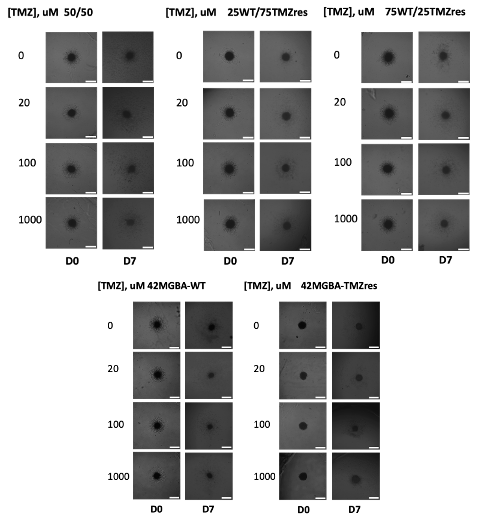

**Figure S2. Raw images of spheroids encapsulated into GelMA matrix and treated with single-dose TMZ.** Brightfield images capture radial spread of GBM cells into the surrounding GelMA hydrogel matrix. Scale bar: 500 µm.

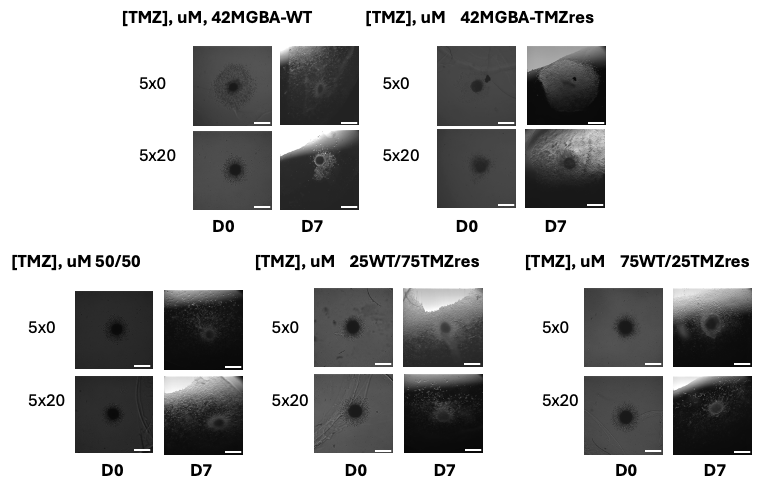

**Figure S2. Raw images of spheroids encapsulated into GelMA matrix and treated with metronomic 5-dose TMZ.** Brightfield images capture radial spread of GBM cells into the surrounding GelMA hydrogel matrix. Scale bar: 500 µm.
